# Malaria shaped human spatial organisation for the last 74 thousand years

**DOI:** 10.1101/2025.06.04.657870

**Authors:** Margherita Colucci, Michela Leonardi, James Blinkhorn, Seth R. Irish, Cecilia Padilla-Iglesias, Stefanie Kaboth-Bar, William D. Gosling, Robert W. Snow, Andrea Manica, Eleanor M.L. Scerri

**Affiliations:** Human Palaeosystems Group, Max-Planck Institute of Geonanthropology, Jena, Germany; Evolutionary Ecology Group, Dept. of Zoology, University of Cambridge, Cambridge, United Kingdom; Natural History Museum, London. Evolutionary Ecology Group, Department of Zoology, University of Cambridge, Cambridge, United Kingdom; Department of Archaeology, Classics and Egyptology, University of Liverpool, United Kingdom; Global Malaria Programme, World Health Organization Avenue Appia 20, Geneva 1211, Switzerland. Swiss Tropical and Public Health Institute (Swiss TPH), Kreuzstrasse 2, 4123, Allschwil, Switzerland; Emmanuel College, University of Cambridge, Cambridge, United Kingdom; Institute of Geological Sciences, Freie Universität Berlin, Berlin, Germany; Institute of Biodiversity & Ecosystem Dynamics, University of Amsterdam, Amsterdam, the Netherlands; Kenya Medical Research Institute (KEMRI)-Wellcome Trust Research Programme, Nairobi, Kenya; Centre for Tropical Medicine and Global Health, Nuffield Department of Clinical Medicine, University of Oxford, Oxford, United Kingdom; Department of Classics and Archaeology, University of Malta, Msida, Malta; Department of Prehistoric Archaeology, University of Cologne, Cologne, Germany

**Keywords:** Malaria, sickle cell anaemia, ancient pathogens, human expansion, *Homo sapiens*, selection, Pleistocene, Species Distribution Models, paleoclimatic reconstructions

## Abstract

The mechanisms driving the spatial organisation of early human societies in Africa are typically addressed through climate variables ^1-3^. However, genetic and archaeological studies have also suggested diseases as a major source of selection in the Pleistocene. Here, we explore whether *P. falciparum*-induced malaria, a major world disease, drove habitat choice in human societies between 74 and 5 thousand years ago (kya). Using species distribution models of three main mosquito complexes with palaeoclimatic reconstructions and combining the results with epidemiological information, we estimated an index of malaria transmission risk in sub-Saharan Africa through time. We then correlated it with an independent reconstruction of the human niche over the same time period and region. Our results show that humans strongly avoided or were unsuccessful in potential malaria hotspots. The effects of these choices shaped human demography for the last 74 kya, and likely much earlier, by fragmenting human societies over time and contributing to the formation of modern population structure. Our results highlight the importance of considering disease distributions when modelling estimates of past human demography, demonstrating that factors beyond climate underlay patterns of human habitat choice, exchange, and dispersal.

**Sentence summary:** Malaria shaped human habitat choice, exchange, and dispersal since the late Pleistocene in sub-Saharan Africa.

## Introduction

Converging evidence demonstrates that our species, *Homo sapiens* did not have one single birthplace in Africa ^4-6^. Instead, the earliest members of our species were divided into small populations that spread across much of the African continent, presenting a dramatically different scenario to long-held perceptions of a single centre of endemism ^4-6^ (Supplementary Information 1). This paradigm shift implicates more than one region and environment in Africa at the root of our species ^6^, and a recognition that humans occupy a “generalist specialist” niche ^7^. Evidence demonstrating that humans, as a species, occupied a wide range of environments from an early stage, while being highly locally adapted at sub-populations level is increasingly apparent (e.g., ^8^). As a result, considerable focus has been given to identify the climate mechanisms of population spread/isolation and ecological adaptation in this context (e.g., ^1-3,9^). However, it was not just humans who adapted to different regions and environments, their pathogens did too. In the current race to elucidate processes of human adaptation and spread, there has been little focus on the link between climate and disease, and the consequent impact on human selective processes ^10-12^. What was the burden of disease in the earliest periods of our species’ prehistory? How did diseases impact human behaviour and demography? How did these factors interact and affected the mixings and dispersals that cumulatively shaped the course of human evolution and, ultimately, the history of all contemporary populations?

Here, we aim to address these questions by studying the impact of diseases in the human past. In particular, we explore how malaria shaped the history of our species in Sub-Saharan Africa between 74 thousand and 5 thousand years ago (kya). This period spans significant demographic expansions within and out of Africa by hunter-gatherer populations ^4,13-16^; up until the time shortly before the expansion of farming lifeways ^17-20^.

Malaria is a major world disease that today presents a global health problem, with 263 million cases annually ^21^. Critically, genetic studies also indicate that malaria was a major problem both in recent prehistory ^22^, and also in the Pleistocene, with mutations relating to sickle cell anaemia emerging in response to malaria between 25-22 kya in Africa (^12^; see SI 2). Archaeological studies have also identified earlier, indirect evidence for the measures humans took to avoid exposure to the vectors of disease, for example, by topping plant bedding with aromatic leaves containing insecticidal and larvicidal chemicals ^23^. Other data may suggest an avoidance of certain localities. For example, the absence of sites near major North African rivers during peak periods of the Last Interglacial (∼125-71 kya), may potentially indicate the avoidance of swampy regions where mosquitos (and other parasites and vectors) thrived ^24^.

Given the indications from genetic archaeological data that malaria may have shaped early human settlement patterns in sub-Saharan Africa, we reconstructed its distribution over time. Reconstructing past disease incidence and its effects on humans is an endeavour that has typically been challenging due to the limited direct evidence from such remote times (see SI 2). We overcame the lack of direct evidence by first quantifying the niche of malaria’s main vectors which, in turn, allowed us to reconstruct their potential spatial distribution for a given point in time based on the climatic conditions. This is done by applying Species Distribution Models (SDMs) ^25^ based on present-day occurrences and location-specific environmental and climatic variables, and then projecting their ranges back in the past. By combining the obtained vectors’ habitat suitability with epidemiological information (see Methods), we then calculated an index of potential risk of malaria transmission, defined as “malaria stability index” ^26^. This index expresses the potential stability of transmission of malaria considering the environmental and habitat conditions favourable to its presence, therefore, quantifying an overall potential risk of transmission. We note that a high stability index does not imply the presence of malaria, but rather defines its potential impact if it was present. In this way, we inferred and mapped the potential stability of malaria through time. By incorporating species ecology, environmental and climatic changes and pathogen stability (i.e., incubation period), we were thus able to map through space and time the areas where malaria had the potential of impacting humans. We then used independent reconstructions of the suitable range for humans (human niche) based on archaeological sites ^13,27^ to track the human expansion across the landscape (see SI 3). Comparing these two reconstructions (potential malaria risk and human ranges) allowed us to infer and quantify the impact of malaria on human demography and dispersal. Our reconstructions show that human behaviour has been substantially shaped by the presence of malaria.

## Results

### Spatial Distribution Models of the studied Anopheles vectors

The use of Species Distribution Models (SDMs) enables us to reconstruct the realised niche of the studied vector species by linking its known occurrences to environmental and climatic variables at their locations to retrieve the potential distribution over a whole area ^25,28^. Anthropogenic land use provides an additional potential driver of the distribution of *Anopheles* mosquitoes (and thus of malaria). To capture the changes in local habitat due to land use change, we use Leaf Area Index (LAI) as a quantitative proxy of the structure of vegetation in both natural and anthropogenic landscapes. LAI was estimated and adjusted for the present by combining estimates based on natural vegetation as predicted from climatic variables with the proportion of habitat converted to crops, pasture and grazing land at any given location (see SI 3). We did not consider human population density, as we wanted to build models that would focus on the climatic drivers of mosquito distributions to be able to project back in the past when humans were hunter-gatherers (and thus their densities were not comparable to current levels) ^29^. Since our SDM focuses on presence over a large geographic area rather than density of mosquitos, this omission should not affect the SDM much, as the presence of a species across the continent should be driven more by the climate, whilst it is its local abundance that will be affected by other factors such as the density of the hosts.

We performed SDM for three selected malaria vector species to include one complex (*An. gambiae* complex), and subsetting two of its salt-water breeding species as a separate group (*An. melas* and *An. merus*), and one group (*An. funestus* group; see Methods). We combined the species observation data with environmental predictors in order to map habitat suitability (from low to high) across the landscape (exemplified in Fig. 1a, c and e) for the whole period analysed (Fig. 1b, d, f). The best models were selected among the predictions in an ensemble ^30^ with a minimum threshold of 0.7 for the Maximum True Skill Statistics (TSS). The obtained vector distributions in the present, reconstructed with these selected variables and map resolution, are coherent with presence data and previous SDM reconstructions ^31^. This latter coherence between our SDMs, which focus on climate, and previous work, which also included human population density, validates our logic that the geographic distribution of mosquitos is mostly driven by climate. We identified the coastal distribution for species such as *An. melas* and *An. merus* (Fig. 1e-f), and highly suitable areas in West and Central Africa for the rest of *An. gambiae* complex species (Fig. 1c-d).

**Figure 1.**
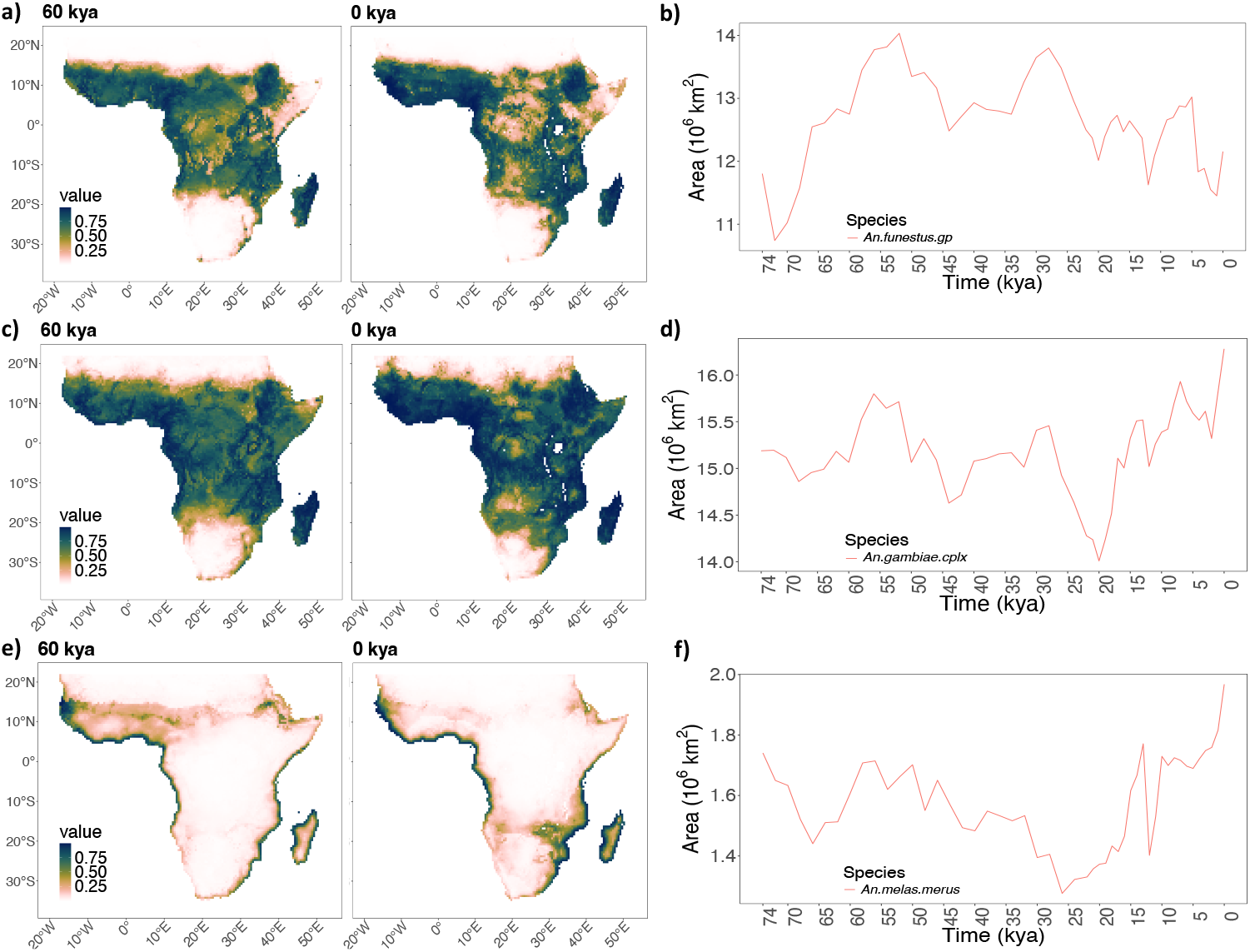
Species distribution models showing the changes in distribution ranges through time. a) SDMs of *An. funestus* group at 60 KYA and in the present (0 KYA) with (b) range of core areas (0.95%) of distribution through time; c) SDMs of *An. gambiae* complex at 60 KYA and in the present (0 KYA) with (d) range of core areas (0.95%) of distribution through time; e) SDMs of *An. melas* and *An. Merus* at 60 KYA and in the present (0 KYA) with (f) range of core areas (0.95%) of distribution through time. Note that the different y-axis in b, d and f reflect the different extent (area) occupied by each species.

### Potential malaria risk through time

We combined the effects of the presence of multiple mosquito species to generate a “malaria stability index” ^26^ representing the potential risk of malaria transmission as an infectious disease at a given location. This potential risk of malaria reflects the stability of transmission (i.e., consistent transmission throughout the year) considering the favourable (ecological) conditions to malaria, conceptually equivalent to the modelling of habitat suitability of a species (e.g., the mosquitoes ecological niche using SDM). This can indicate the ecological conditions linked to high risk of malaria and, consequently, malaria’s potential ranges. Thus, current sedentary lifestyles and the presence of people do not have an impact on the potential ranges, but affect the incidence of malaria, increasing the risk of transmission. The malaria stability index, therefore, expresses the stability of transmission and combines the impact of each vector present based on its ecology and its contribution to the persistence of *P. falciparum* (factors that consistently and strongly impact the stability of malaria transmission), and it has been shown to correlate well with the incidence of the disease ^26^. We computed the malaria stability index over the last 74 kya across sub-Saharan Africa.

The pattern that emerges shows a clear increment of malaria stability over time across the landscape (Fig. 2). The first visible peak of malaria corresponded with the main “Out of Africa” expansion at about 60-50 kya, and is in line with the suggestion that malaria travelled with humans in their migrations outside the African continent at this time ^32^. Then, the major peak occurred shortly after the Last Glacial Maximum, at around 13 kya, showing that an increase in the extent of areas with high malaria stability was taking place well before agricultural practices began to emerge around 8 kya. This result supports the finding by Laval and colleagues ^12^ that selection pressure of malaria predated food production. We note that our results do not conflict with the suggestion that sedentary lifestyles and increased population sizes had independent effects on local incidence of malaria. As we highlighted before, the obtained malaria stability index expresses the potential risk of malaria and quantifies its range.

**Figure 2.**
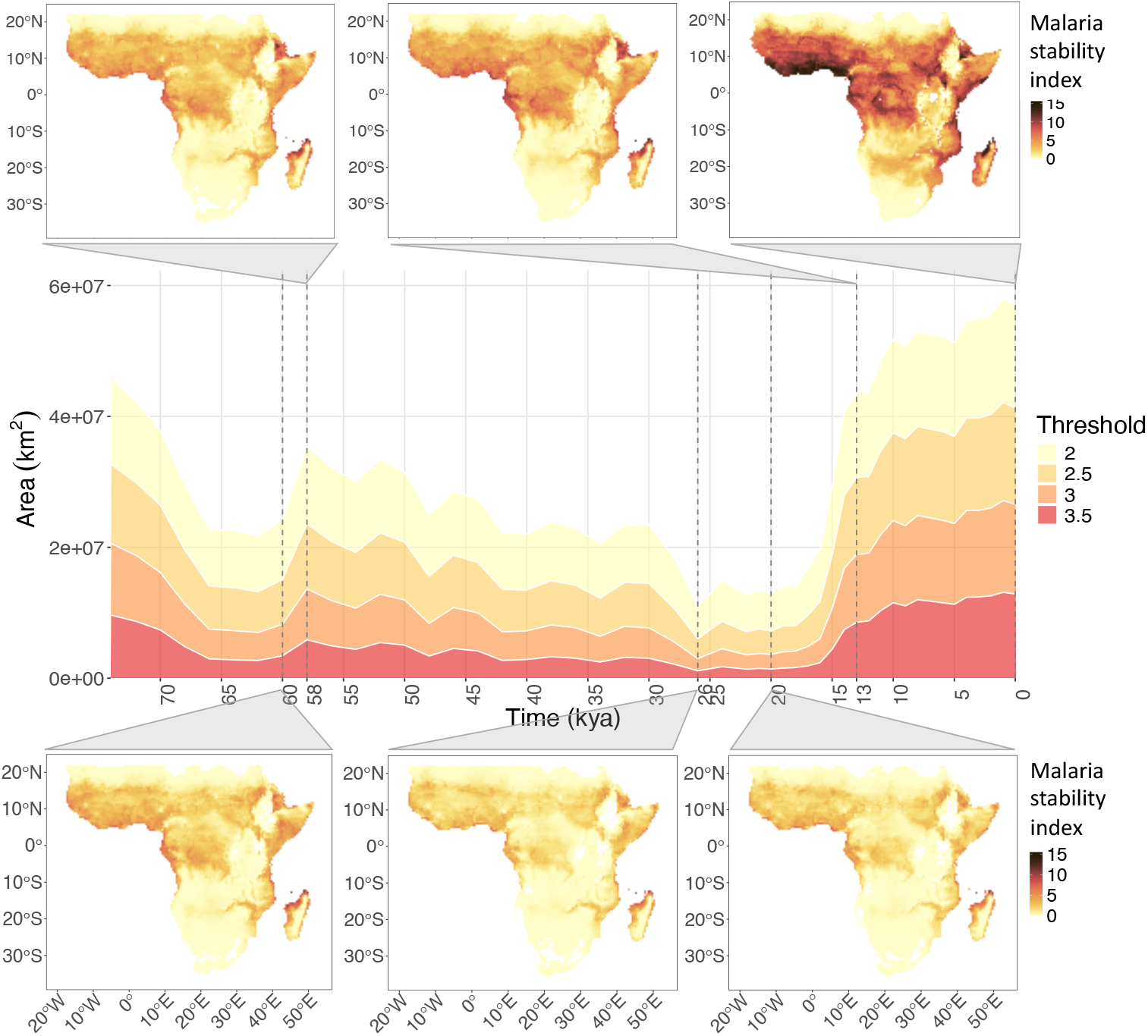
Malaria stability index through time, considering an environment impacted by land use. The areas in the map (km^2^) that have an index value above a set of arbitrary thresholds (i.e., high malaria levels) are on the y-axis, while time (thousand years ago, KYA) is on the x-axis. The same pattern can be seen independently from the choice of threshold. Six different time steps are highlighted: at 60 kya, 26 kya and at 20 kya we see a decrease in stability of overall malaria, while at 58 kya, 13 kya and present (0) we see an increase.

### Malaria impact on human range expansion

We then tested the impact of potential malaria risk on human settlement patterns by comparing the changes in our malaria stability index reconstructions across sub-Saharan Africa with independent reconstructions of the potential distribution of hunter-gatherers over the same period and area (based on ^27^, an extended version of ^13^, see SI), considering a time frame from 74 to 5 kya to focus only on the hunter-gatherer population. These reconstructions of hunter-gatherers’ distribution were obtained using SDMs to reconstruct the ecological niche of humans by modelling the relationship between climatic variables and the distribution of archaeological sites (see Methods). These occurrences were based on a curated pan-African dataset of archaeological sites (based on ^13^, and expanded to 5 kya ^27^), and allowed the niche to change through time ^33^. It is worth highlighting that this reconstruction is completely independent from our analyses on malaria, but, because it uses the same palaeoclimatic data ^34,35^, we can compare the two reconstructions.

The pattern that emerges reveals a negative relationship between areas of high malaria stability and suitability for *Homo sapiens*. This indicates that human expansion within Africa was significantly impacted by the presence of malaria and that humans likely avoided areas with a high potential risk of malaria transmission through time (Fig. 3, see Fig. S4 for more time steps). In Figure 3a, the regions most likely inhabited by humans (core areas, see Methods) are superimposed (black outlines) onto the map of the potential risk of malaria. The figure shows how areas of low stability of malaria transmission were consistently more suitable to human inhabitation. Through time, this created potential corridors for human movement and expansion as well as pockets of isolated human groups. For example, it is possible to notice that areas between the Saharan and Ethiopian habitable zones underwent cycles of disconnection (e.g., at 54 and again at 8 kya in Fig. 3) and connection (e.g., at 16 kya in Fig. 3).

**Figure 3.**
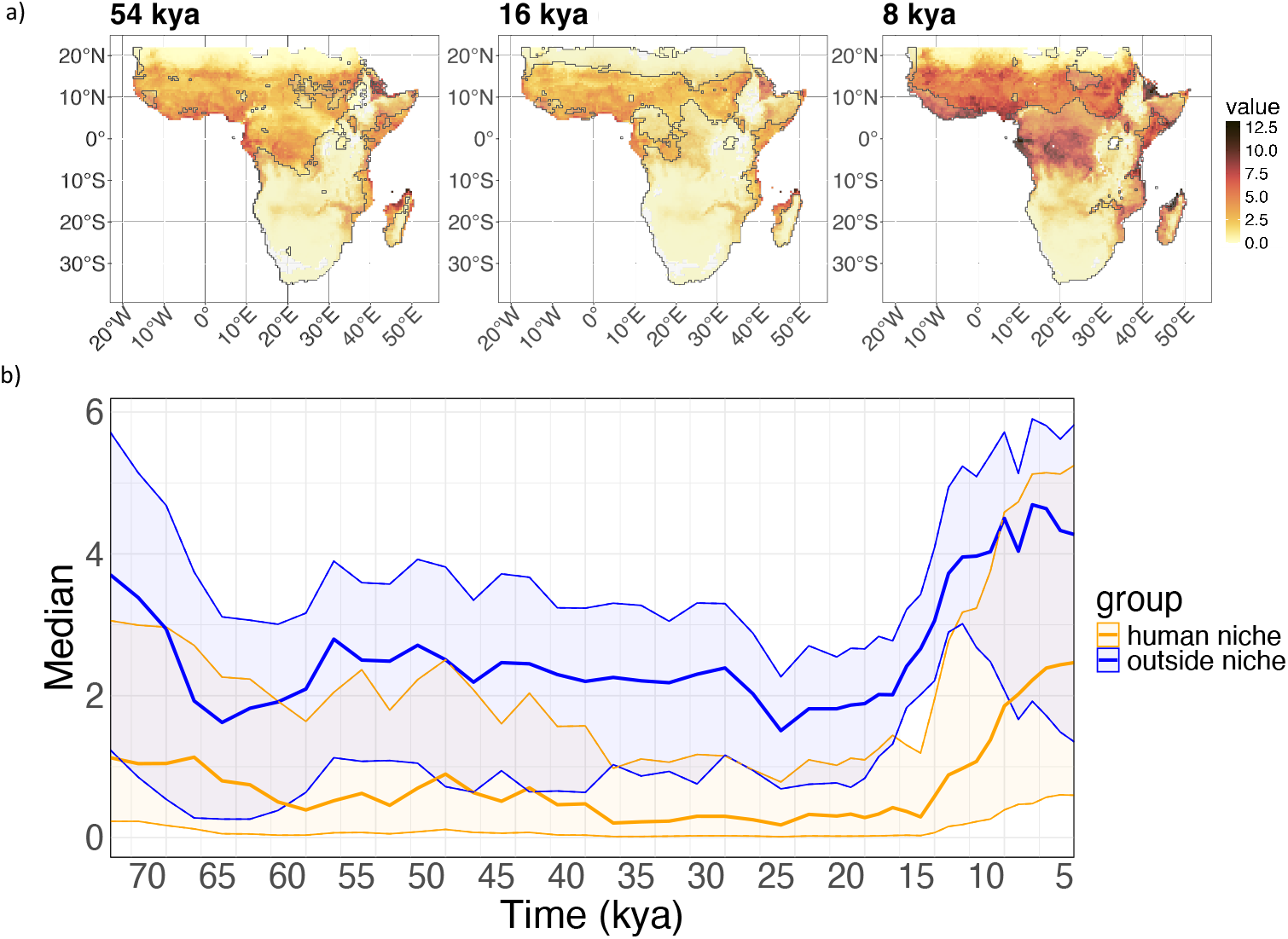
Comparing the extent of human niche and malaria stability index through time. a) Extent of the human niche (outlined in black) against the map of malaria stability index at 54, 16 and 8 kya; b) median of level of malaria stability index in the area of human range (dark orange line) and outside the area of human range (dark blue line), including the uncertainty (interquartile, colour in transparency around the darker lines that shows median values). We can see that the level of malaria in the human niche is consistently lower than the areas avoided by humans.

To further explore the pattern, we estimated the level of malaria stability index (median and interquartile) across the landscape, dividing the map in (core) “human niche” (based on the core area that would include 95% of archaeological sites) versus “outside the niche”. We can see that the level of malaria stability index was much lower in the areas suitable for humans than in the areas outside the human range, and this is consistent through time (Fig. 3b). The human niche was characterised by consistently low values (Fig. 3b and, for example, 54 kya in Fig. 3a) from 74 kya until around 13 kya. It is also possible to notice that from around 14-13 kya, the overlap between malarious areas and the periphery of human ranges increased, meaning that humans were possibly venturing in areas with higher malaria risk they were not reaching before. By 10 kya, we can see increased overlap, reaching the highest values especially in West Africa (see 8 kya in Fig. 3a) where we expect the initial spread of the resistance mutation sickle cell anaemia.

## Discussion

Our results show that the potential risk of malaria transmission shaped the spatial organisation of human groups at least over the last 74 kya, structuring populations into different regions of sub-Saharan Africa, and creating habitat “islands” that isolated some local populations from others. This increased overlap between high malaria risk areas and human ranges becomes more evident after 15-14 kya. It is plausible that this pattern reflects the spread of resistance mutations such as sickle cell anaemia in West and Central Africa ^36^, supported by the timings of selection suggested by ^12^, and the hypothesis that links sickle cell to Bantu population in these areas ^37-39^. Furthermore, our results show that malaria was already at an extremely high level around 13 kya, before the suggested advent of agro-pastoral lifestyle around 8-7 kya. Our results, therefore, show that, due to climate conditions, malaria had already a potential major driver of selection before food production. These results also caution against making assumptions regarding links between Neolithisation and the origins of several infectious diseases (e.g., ^40-43^). How and why pathogens emerged and spread, and impacted human populations beyond health is clearly a critical dimension of the human past that is still largely unexplored.

Until now, the lack of availability of ancient genomes for direct evidence and refined dating of malaria in the Pleistocene has been thought to present a challenge by limiting studies to historical and modern data from *Plasmodium* ^22,44,45^ and from living people ^12^. This study provides proof of concept that we can explore disease burden in the earliest periods of human prehistory without direct evidence of human-pathogen interactions. It is also instructive that climate alone can potentially determine disease presence without complex anthropogenic habitat intervention, or major changes in subsistence economy.

This could be the case for many diseases. Our methods and approach provide a way to understand disease burden in the past and how this may have impacted the spatial organisation of human groups, their patterns of dispersal, and their degree of contact/isolation, representing an entirely new dimension of knowledge for understanding the dynamics shaping the formation of our species.

## Methods

### Anopheles species

We studied three *Anopheles* mosquito groups that are major vectors of malaria: *An. gambiae* complex, *An. melas* and *An. merus* (considered as an independent group being a subset of *An. gambiae* complex with specific saltwater-breeding habits), and *An. funestus* group. It is important to note that differentiation of sibling species within the *gambiae* complex was not possible until the 1990s, or earlier through cross-mating (e.g., differences between *An. melas* and *gambiae*, or between *An. gambiae* and *An. arabiensis* in the 1960s). Different *Anopheles* species show different breeding periods, feeding patterns, survival rates, and, therefore, different distribution and competence ^26,31^, strongly determined by the climate and environment ^46^. It is also important to consider the interactions between vector species: when a habitat is invaded, other vectors may be displaced. A competent vector (i.e., a vector that is anthropophilic, more abundant than other *Anopheles* species and that frequently contains sporozoites) is defined as “dominant” (or primary) ^26^. Kiszewski co-workers’ model (2004) recognises a different occurrence of dominant vectors during different seasons within a region, considering only the contribution of one dominant vector as the most significant in determining endemicity in the region, while ignoring secondary vectors ^26^. This information and expert contribution guided this study in the choice of species to include in the model, creating a curated dataset that ensured correct taxonomic identification.

Among the *Anopheles* species that are known to inhabit sub-Saharan Africa, we selected dominant vectors and relevant complexes to cover different geographical distributions, biome and host preferences (expert information; ^31,47^). See Table S1 for the list of species and sibling groups covered.

### Summary of species distribution map generation

#### Environmental and climatic data

The *Anopheles* mosquito vector species are influenced by climate, which is an important factor for the vector’s spatial distribution range ^48^. Individual sibling complexes show different relationships with climatic factors, especially temperature, humidity/wetness ^31^ and precipitation ^49^. Temperature also affects the duration of sporogony of the parasite in the mosquito ^50^.

The area of study was limited to sub-Saharan Africa including Madagascar. To reconstruct the environment map, we used a dataset (via *pastclim* v 1.2 ^51^) at a native resolution of 0.5° and continuous palaeoclimatic reconstructions from the present with intervals of 1000 years up to 22 kya, then of 2000 years up to 74 kya ^34^. This dataset includes 17 bioclimatic variables: all the BioClim variables excluding BIO15 (in ^34^), Leaf Area Index (LAI), Net Primary Productivity (NPP) and rugosity (a measure of the standard deviation in altitude within a certain area to reflect topography).

To reconstruct both a natural, pristine environment and a modified environment (i.e., impacted by agriculture or pastoralism, here defined as “land use”) we adapted land use variables from the History Database of the Global Environment (HYDE version 3.2, ^52^). We considered cropland, grazing land, and pastures from HYDE v. 3.2 to reflect the type of vegetation in Africa from 10 kya until the present. To define an “open” and “closed” type of vegetation and, therefore, reflecting the type of environments favoured by mosquitoes, the Leaf Area Index (LAI) was used as a proxy. The cropland and pasture variables were converted to LAI (with LAI of 1.7 for both ^53,28^) creating the variable “land use”. Grazing variable from HYDE v. 3.2 was used as a proxy for cattle. Additionally, the distance from the sea (in km) was included as a variable for coastal vector species like *An. merus* and *An. melas*.

### Species Distribution Models

#### Data cleaning and thinning

Datapoints of presences were obtained from ^31,47,54^. Using the R package *tidysdm* ^55^, the data points were thinned, keeping one observation per cell, with a distance of 70 km between each observation.

#### Model fitting

SDMs were performed using the R package *tidysdm* ^55^. The thinned data set was used as presences ^31,47,54^, and three times the number of presences was drawn (20 times) as pseudo-absences keeping a minimum distance of 150 km from the presences.

The variables of interest were chosen considering the proportional overlap of presences’ and pseudo-absences’ distributions over the variable space (*tidysdm::dist_pres_vs_bg()*), where high distance between the two distributions is indicative of non-random use of the area by the species (Table S3). Climatic and environmental variables were further selected to better capture the difference between Western and Eastern Africa at the current map resolution (0.5° x 0.5°), leaving BIO5, BIO6, BIO4, BIO8, BIO16, BIO18, BIO19, NPP, rugosity, LAI, grazing, and distance from the sea. Then, these selected variables were pruned for collinearity (i.e., greatest mean correlation, with a cutoff *r* of 0.8).

The models were run for four different algorithms using *tidysdm*: generalised linear models (GLMs), random forest, generalised boosting method (GBM), and MaxEnt (see SI).

Next, using the *tidymodels* approach ^56^, different models were fitted defining a workflow. Models tuning and evaluation was performed by a spatial block cross-validation scheme ^57^. The data is divided in a 80:20 split (i.e., 4/5 of the splits are used for calibrating the model and the remaining 1/5 for evaluation) by creating 5 folds, and 20 combinations of the hyperparameters were explored for each algorithm (based on ^58^).

Then, we created an ensemble ^30^ using the Maximum True Skill Statistics (TSS) as a metric to choose the best random forest, boosted tree, MaxEnt and GLMs. Members added to the ensemble are fitted to the full training dataset to be used for predictions. The tidysdm function *repeat_ensemble()* was used, based on 20 resampled datasets (20 iterations for each species) to explore further the effects and performance of thinning data and selecting pseudo-absences. With this ensemble, it is possible to make predictions based on the median of the best models setting a minimum TSS threshold of 0.7.

The present predictions were in line with the outputs from other SDM studies ^31^ and ^54^, confirming out choice of variables (including the impact of land use change). We then projected the obtained model into the past up to 74 kya (Figure S3 showing some time steps for some species). The contribution of each variable to the models was also explored (Table S3).

#### Epidemiological information: the malaria stability index

Once the niche of dominant species of mosquitoes was identified through time, we calculated an index for the stability of malaria transmission at each time step, based on ^26^. The index includes *P. falciparum* incubation period, vector’s biting activity (i.e., proportion of bites on humans by the dominant vectors), daily survival rate of the vectors and monthly temperature:

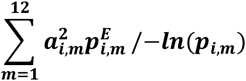

where *E* = 111/*Temperature* ™ 16°C for *P. falciparum*.

Finally, we multiplied the median probability of vectors’ incidences from the obtained SDMs by the malaria stability index to find the areas with the highest incidence rate. For a detailed explanation of the methods and interpretation of the malaria stability index, see the Supplementary material.

#### Correlating the human niche to the malaria stability index

Independent reconstructions of the human niche were obtained from ^13^ and ^27^ (see SI). The reconstructions were based on archaeological sites across Africa, with dating starting from 120 kya, considering five climatic and environmental variables ^59^: leaf area index (LAI), temperature annual range (BIO7), mean temperature of the wettest quarter (BIO8), mean temperature of warmest quarter (BIO10), and precipitation of wettest quarter (BIO16). The niche area was divided into core areas (the smaller area that includes the 90% of the presences) and extended areas (covering 95% and 99%) by ^13^. We chose a level of 95% as the range of humans across the landscape (see ^13^, where “core area” is instead defined as an area encompassing 90% of presences).

Firstly, we checked the spatial overlay of the human core area against the maps of malaria stability index extent across time (see Fig. 3a). Then, we compared the median level of the malaria stability index in the areas identified as core areas for humans against the median level of the malaria stability index in the areas outside the human range, i.e., areas identified as not suitable for human groups (considering upper and lower quantiles of 0.25 and 0.75). Desert areas (unsuitable for both humans and malaria vectors) were still included in this analysis.

## Acknowledgments

RWS is supported as a Wellcome Trust Principal Fellow (#212176) and is grateful for the support of the Wellcome Trust to the Kenya Major Overseas Programme (#203077). M.C. and E.M.L.S. were funded by the Lise Meitner Pan-African Evolution Research Group. ML and AM were funded by the Leverhulme Research Grant RPG-2020-317

## Author contributions

Conceptualization: EMLS, AM

Project Methodology: EMLS, AM, MC, ML

Investigation: MC

Expert advice, methodology: RWS, SI, CIP, JB, SKB, WG

Funding acquisition: EMLS

Project administration: EMLS, AM

Supervision: AM, EMLS

Writing – original draft: MC, AM, EMLS

Writing – review & editing: all authors

## Competing interests

The authors have declared that no competing interests exist.

## Notes

### Competing Interest Statement

The authors have declared no competing interest.

