## Supplementary Information for "Malaria shaped human spatial organisation for the last 74 thousand years"

**SI 1. Human Origins**
**SI 2. The study of past diseases: malaria as a case study**
**SI 3. Methods**

### 1. Human Origins

In the last few years, there has been a profound reconsideration of the character of human evolution, and the origin of our species *Homo sapiens* in Africa. Rather than, as the word "origin" suggests, emerging from a single centre of endemism, the set of cognitive and behavioural traits that define our species emerged through an evolutionary process that played out across much of the African continent. Over time, and through the genetic exchanges between diverse sets of local populations living in different regions of Africa, the set of physical and behavioural features that defines *H. sapiens* began to emerge. This pattern is evidence both through fossils and material culture. Fossils bearing modern traits appear in different African regions from around 300 thousand years ago (kya) [1,2], with some archaic features lasting until relatively recently [3]. Similarly, Middle Stone Age (MSA) material culture, which marks a fundamental re-organisation of technological conceptions, appears with some of the earliest *Homo sapiens* fossils around 300 kya [1,4]. Thought to be the earliest manifestations of modern cognition, the MSA also appears in many different African regions at around the same time [2,5].

Most recently, studies using genetic data from contemporary populations have supported the view that humans are one species with several African roots, bearing out a range of prescient model-based and theoretical studies [2,6,7]. Ragsdale and colleagues (2023) [8] presented a study showing that multiple stem populations contributed to the emergence of our species.

This early population structure is therefore key to understanding the character of modern human origins. However, the mechanisms driving population structure are not well understood. Currently, all that seems clear is that the earliest members of our species lived within a structured metapopulation, whose dynamics shifted over time, with modern-day population structure originating before 300 kya [9-11]. It seems likely that some of the same mechanisms drove this shifting and dynamic population structure. Chief among candidate mechanisms is climate. Africa's fractured palaeoclimates have been well documented, particularly from the Last Interglacial (~130 kya) [12-15]. The ebb and flow of the Sahara Desert [16], "mega" droughts in Central Africa [17], moisture “see-saws” between eastern and western Africa [14] among other examples, all contributed to differential climate dynamics across Africa. As a result, they are also likely to have impacted the dynamics of human habitation, dispersal, and encounters between local groups that underpin genetic exchanges. They may also underpin similar patterns of sub-structure observed in the genetics of other pan-African mammal species [18].

Beyond this, geographic distances, and potentially, culture drove some degree of population structure, as they do today [5]. However, disease has historically not been considered as a factor driving habitat choice and human demography in the African Pleistocene, despite the major influence on human populations’ health.

### 2. The study of past diseases: malaria as a case study

Diseases have shaped human behaviour through time, often in extreme ways, from past epidemics (e.g., plague), to present pandemics (e.g., COVID-19). However, our ability to analyse and quantify the actual disease burden has been limited, especially when considering longer times-scales. For example, recent studies revealed the complex history of hepatitis B virus (HBV), revealing it presence in Neolithic, Bronze Age, as well as Iron Age [19,20] and transmission between ancestors of European and Native American populations around 15-13 kya [19] highlighting the ancient origin of this disease [21] and suggesting the potential burden on the population already in the late Pleistocene. However, no direct evidence of prevalence or effects is available on this ancient virus.

Some vital information can be obtained by the application of indirect methods. For example, genetic analysis of modern (and ancient) human samples enables us to indirectly explore the impact of pathogens on human genomes (and populations). This can be done by examining the selective pressure imposed by the disease as a result of host-pathogens encounters and pathogenic load [22-25]. An example is the impact of malaria on human populations: due to its significant burden on human health, several mutations that offer a level of protection emerged alongside it, making this disease ideal for our study.

Due to its high morbidity and mortality, with around 263 million cases globally [26] (Fig. S1). Malaria is caused by *Plasmodium* parasites, of which five types exist: *Plasmodium falciparum*, *Plasmodium vivax*, *Plasmodium malariae*, *Plasmodium ovale*, and the zoonotic/simian *Plasmodium knowlesi*. Among these, *P. falciparum* is the most virulent of human malaria parasites [27]. The extensive distribution of these parasites and the burden of the disease placed one of the highest selective pressures on the human genome, leading to the emergence of resistance mutation (i.e., mutations that offer a level of protection), such as sickle cell anaemia [28], α and β-globin mutations in thalassemia, Duffy blood group variants [29], tumour necrosis factor (TNF), and glucose-6-phosphate dehydrogenase (G6PD) [30-32].

In this paper, we focus on *P. falciparum* and haemoglobin $\beta^s$ sickle mutation, which has been reported with high frequency in areas of high malaria density, particularly in sub-Saharan Africa [33,34]. While the homozygous form of this mutation can lead to sickle-cell disease and potentially death, the heterozygous form offers higher protection against malaria [35,36]. Because of its distribution and malaria-resistance effect, the emergence of $\beta^s$ sickle mutation in humans has been studied to better understand the history of malaria itself.

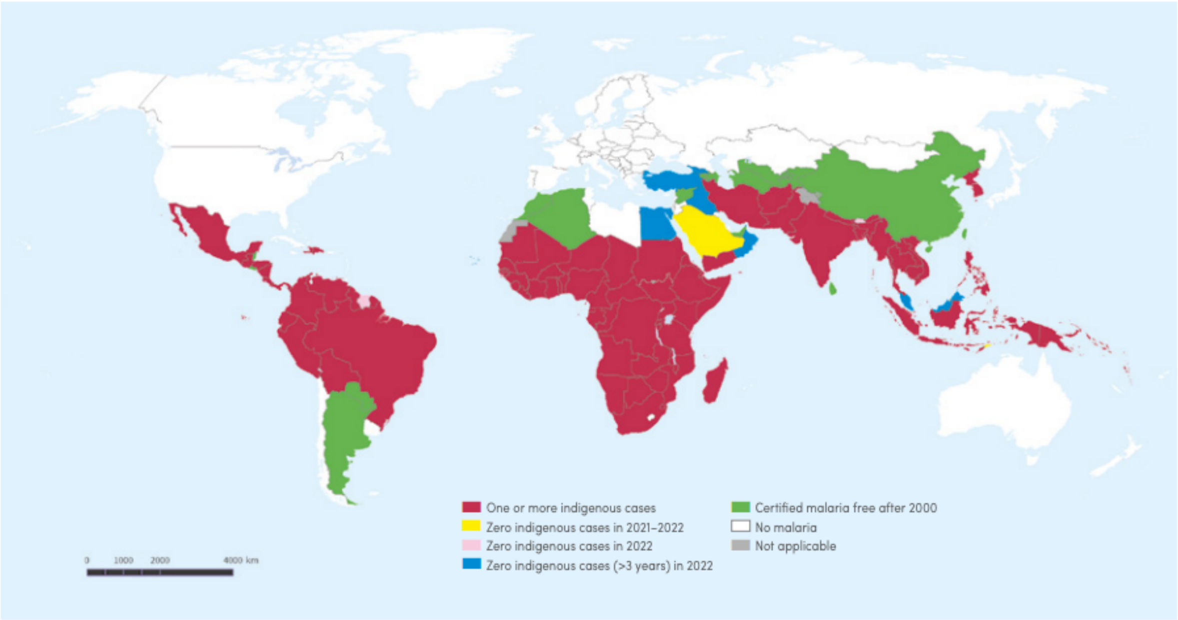


**Figure S1: Global map of indigenous cases of malaria in 2022.** This map shows the extent of malaria in 2000 and the cases in 2022. Source: World Health Organization database [26].

### Theories on the origin of malaria (*P. falciparum*)

Despite this attention, very little is known about malaria's epidemiological and evolutionary past and, therefore, about the plethora of conditions linked to its historical risk that are relevant to current challenges and prevention efforts [37].

There are various hypotheses, considering both a multicentric model, with five independent occurrences spanning a time range of a few thousand years, and a unicentric model, with an older, single occurrence [38]. These hypotheses consider central [28,39] and west Africa as points of origin [33,40,41]. These areas are further explored in this paper.

Traditionally, malaria is linked to the emergence of slash-and-burn agriculture in the past 10-8 kya [28]. Authors have highlighted fundamental factors such as a more favourable climate and environment in Africa (about 12-7 kya) for plant and animal domestication (about 10-8 kya in the Sahara and northeast Africa), for human settlements as well as malaria vector species [30]. Considering the emergence of the original sickle haplotype, a time frame predating the Bantu expansions, around 7300 ya, was proposed [38]. However, there are various lines of evidence that nonetheless backtrack its presence.

Considering the age of *P. falciparum*, it was demonstrated that the plasmodium existed before agriculture (90% credibility intervals), and it was linked to the movements of the human host from sub-Saharan Africa, following humans during an expansion out of Africa around 60–50 kya ago [42].

Another line of evidence comes in the form of genetics selective pressure due to malaria disease burden. Laval and colleagues [24] tracked sickle cell mutation to date the start of selective pressure by the most virulent form of malaria, transmitted by *Plasmodium falciparum*. Focusing on rainforest hunter-gatherers and agriculturalist groups, they identified 25-22 kya as the start of sickle cell anaemia distribution/the rise of the sickle cell mutation. This study highlights the ancient origin of malaria, breaking the usual narrative that links the emergence of diseases to the emergence of food production (e.g., [43-46]). The frequencies of sickle cell used in [24] are inferred from modern individuals from publicly available resources of global human genetic diversity (1000 Genomes Project [47] and HapMap [48]). However, it also offers a wide range of confidence intervals on dates: the question on when malaria, as a significant disease developed, is still open.

With our methods, we propose a reconstruction of the potential risk of malaria in sub-Saharan Africa since the late Pleistocene. Our reconstructions corroborate Laval et al. [24] findings, and allow us to infer and, for the first time, quantify how human behaviour is heavily shaped by the presence of a disease.

**The focus on malaria vectors**

Understanding malaria by studying its vectors is crucial [49,50]. Malaria is transmitted by mosquitoes species in the *Anopheles* genus, which includes around 400 species, with about 70 of relevance for human health [51]. The diversity shown among *Anopheles* species highlights the importance of identifying and mapping the most significant malaria vectors.

Particularly in Africa, it is possible to observe significant variability in the geographic distribution of sibling species. For instance, *An. gambiae* has a wide range, expanding particularly in West Africa, while other species, like *An. melas* and *An. merus,* favour coastal areas (i.e., saltwater breading), but can be found further inland too [49]. *An. funestus* has extended ranges to the point of being described as "ubiquitous" [49]. Therefore, a better understanding of the ecological factors that favour anopheline species to implement the use of ecology-based models is necessary for understanding and predicting malaria transmission and distribution [52].

**The question of environmental and anthropogenic changes**

As described in the previous section, plasmodium survival and malaria vector distribution are profoundly affected by favourable environmental and climatic conditions. Currently, malaria incidence is determined by population density and linked urbanisation, with significant impact from climatic and environmental factors such as temperature and water availability. Also, the increase of open habitat, such as the conversion of natural land cover to cultivated land, favours an increase in the range of the mosquito vector [53].

In this paper, we considered the environmental conditions prior to possible human expansion and intervention, and the subsequent changes through time from the Late Pleistocene.

Gosling et al. (2022) highlight that the available records (from western and central Africa, Lake Bosumtwi and Lake Bambili) show significant vegetation change (from forest to savannah, with a decline of tree line) around 300–50 kya [15]. Despite being generally associated with this type of open landscape, humans and evidence of human activity can be found in a variety of different environments, also attested by a change in technology and tools to possibly accommodate new environmental challenges [15].

### 3. Methods

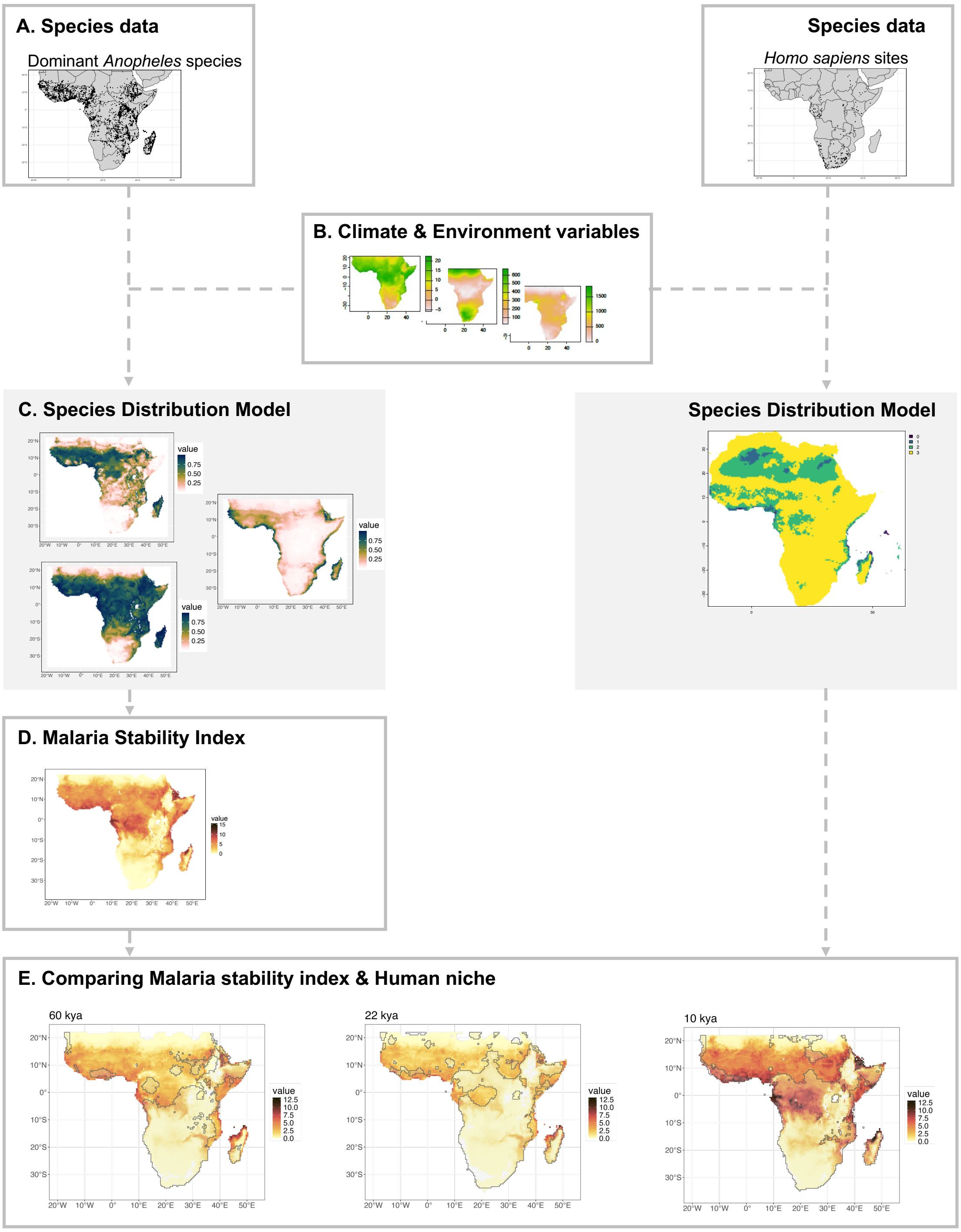


**Figure S2. Overview of the methods and steps to model malaria through time.** (A) starting with the species data points (observations in the present for each mosquito species or sites of *Homo sapiens* through time), we then included climatic and environmental reconstructions (B) to create independent reconstructions of the species ranges (Species Distribution Models) ( C). The mosquito SDMs are then summarised and, including epidemiological information, (D) the malaria stability index is calculated. Finally, (E) we checked the spatial overlay of the human niche against the maps of malaria stability index extent across time.

To explore the extent of malaria spread in the past and its influence on human population ranges, we reconstructed the environmental niche of its sub-Saharan mosquito vectors, the inferred the potential risk of malaria (i.e., malaria stability index), and we compared the potential risk of malaria against the potential range of *Homo sapiens*.

The workflow to reconstruct past malaria distribution includes three main steps (see Fig. S2):

- Species Distribution Models (SDMs) based on (A) observed presences of mosquito vectors and, independently, on archaeological sites associated with *H. sapiens* presence across time and space, reconstructing their distribution, considering climate and environmental models (steps B-C)
- Calculating the malaria stability index based on climatic and epidemiological information on the plasmodium and the vectors (step D);
- Then, comparing the obtained independent reconstruction (SDMs, step E) of the human habitat range across Africa against the obtained range of potential risk of malaria.

These steps will be described in detail in the next paragraphs, starting with the description of data used.

#### *Anopheles* species

A competent vector (i.e., a vector that is anthropophilic, more abundant than other anophelines species and that regularly harbours sporozoites) is defined as "dominant" (or primary) [54]. Kiszewski co-workers' model (2004) [54] recognises occurrences of different dominant vectors during different seasons within a region, considering only the contribution of one dominant vector as the most significant in determining endemicity in the region, while ignoring secondary vectors. This information guided this study in the choice of species to include in the model. We studied three *Anopheles* mosquito groups that are dominant vectors of *P. falciparum*: *An. gambiae* complex, *An. melas*, *An. merus*, and *An. funestus* group. Different *Anopheles* species show different breeding periods, feeding patterns, survival rates, and, therefore, different distribution and competence [54,55], strongly determined by the climate and environment [56]. Their occurrence and abundance are also influenced by interactions between vector species: when a habitat is invaded, other vectors may be displaced. This is the case for *An. funestus* and *An. gambiae* complex species, as they interact across the season maintaining a certain level of malaria yearly [54].

Among the *Anopheles* species that are known to inhabit sub-Saharan Africa, we selected dominant vectors and relevant sibling complexes to cover different geographical distributions and biomes (expert information; [55,57]). These species have been demonstrated to consistently transmit malaria in the studied area [54]. *An. gambiae, An. coluzzii*, and *An. arabiensis* are three vectors (here included in the *An. gambiae* species complex as in historical records) that show the most extensive ranges across sub-Saharan Africa, with some possible natural barriers (e.g., *An. gambiae* is limited by Rift Valley complex [58,59], *An. coluzzii* by the Congo Basin tropical rainforest, and *An. arabiensis* by the Indian Ocean) hindering possible migration [59]. See Table S1 for the list of species and sibling groups covered.

*Data sources*

Presences were obtained from [49,55] and expert information, creating a curated dataset based on taxonomic identification and able to cover a wide range of species-specific habitats. The observations were limited to sub-Saharan Africa, including Madagascar, but excluding smaller islands such as Comoros and Mayotte.

### Summary of species distribution map generation

### Environment and climate reconstructions

The *Anopheles* mosquito vectors' spatial distribution ranges are influenced by climate [60]. Individual vectors show different relationships with climatic factors, especially temperature and humidity/wetness [55] and precipitation [51]. Wet and dry cycles maintain levels of malaria transmission through interactions between dry-season and wet-season vectors [54]. Stable levels of heath favours stable presence of mosquitoes (e.g., savannah areas in west and central Africa) [54]. Temperature also affects the duration of sporogony of the parasite in the mosquito [61]. Topography is also considered an important factor alongside climate and anthropogenic intervention [53].

The area of study was limited to sub-Saharan Africa including Madagascar, following the distribution of sickle cell anaemia and *P. falciparum*. To reconstruct the environment map, we used a dataset at a native resolution of 0.5° x 0.5° and time step reconstructions from the present with intervals of 1000 years up to 22 kya, then of 2000 years up to 74 kya [62]. This dataset includes 19 bioclimatic variables: all the BioClim variables (excluding BIO15) [63], Leaf Area Index (LAI, a measure of canopy foliage used as a proxy for vegetation), Net Primary productivity (npp) and rugosity (a measure of the standard deviation in altitude within a certain area to reflect topography).

To reconstruct both a natural, "pristine" environment and a "modified" environment (i.e., impacted by conversion of land to crop, pasture and grazing land with opening of the vegetation, here described as "land use" and "land use with grazing") we adapted land use variables (i.e., cropland, pasture, grazing) from the History Database of the Global Environment (HYDE version 3.2, [64]). We considered cropland, grazing land, and pastures from HYDE v. 3.2 to reflect the type of vegetation in Africa from 10 kya until the present. The effect of land use was modelled by using the Leaf Area Index (LAI) thus defining an "open" and "closed" type of vegetation and, therefore, as proxy for the type of environments favoured by mosquitoes. The cropland and pasture variables were converted to LAI (with LAI of 1.7 for both) creating the variable "land use" considering the proportion of LAI corresponding to an agricultural environment in Africa (which we considered to be closer to a grassland landscape as calculated in [65,66] rather than intensive cropland from world-wide measurements [65,66]). Grazing variable from HYDE v. 3.2 was used as a proxy for cattle. Additionally, the distance from the sea (in km) was included as a variable for coastal vector species like *An. merus* and *An. melas*.

**Species Distribution Models**

The use of Species Distribution Models (SDM) enables to reconstruct the realised niche of the studied species by mapping its occurrences at known locations to the environmental variables of its habitat [66,67]. This produces a model of the possible distribution of the species across the landscape informed by the most suitable climatic conditions.

**Model fitting**

SDMs were performed using the R package *tidysdm* [68]. The thinned data set was used as presences (one observation per cell, with a distance of 70 km between each observation), and three times the number of presences was drawn (20 iterations) as pseudo-absences keeping a distance of 120 km from the presences.

Climatic and environmental variables were selected to better capture the difference between West-East Africa at the current map resolution and to reveal a difference in distribution between presences' and pseudo-absences' distributions over the variable space

[68,69], leaving BIO5, BIO6, BIO4, BIO8, BIO16, BIO18, BIO19, NPP, rugosity, LAI, grazing, and sea distance. Then, as environmental variables often show collinearity (which can affect models such as generalised linear models), these selected variables were pruned for collinearity (i.e., greatest mean correlation, with a cutoff *r* of 0.8).

The models were run for four different algorithms (ensemble modelling framework) as implemented in *tidysdm*: generalised linear models (GLMs), random forest, generalised boosting method (GBM), and maxent [70].

Next, using the *tidymodels* approach [71], different models were fitted defining a workflow. Models tuning and evaluation was performed by a spatial block cross-validation scheme [72]. The data is divided in a 80:20 split (i.e., 4/5 of the splits are used for calibrating the model and the remaining 1/5 for evaluation) by creating 5 folds and 20 combinations of the hyperparameters (based on [73]).

Then, we created an ensemble combining the four selected machine learning algorithms using the Maximum True Skill Statistics (TSS) as a metric to choose the best random forest and boosted tree. Members added to the ensemble are fitted to the full training dataset to be used for predictions. The *tidysdm* "repeated ensemble" option was used, creating 20 datasets (20 iterations for each species) to explore further the effects and performance of thinning data and selecting pseudo-absences. With this ensemble, it is possible to make predictions based on the median of the best models setting a minimum threshold of 0.7 for TSS.

The present predictions were in line with the outputs from [49,55]. We then projected the obtained model into the past up to 74 kya (Figure S3 showing three time steps for each species). The contribution of each variable to the models was also explored (Table S3).

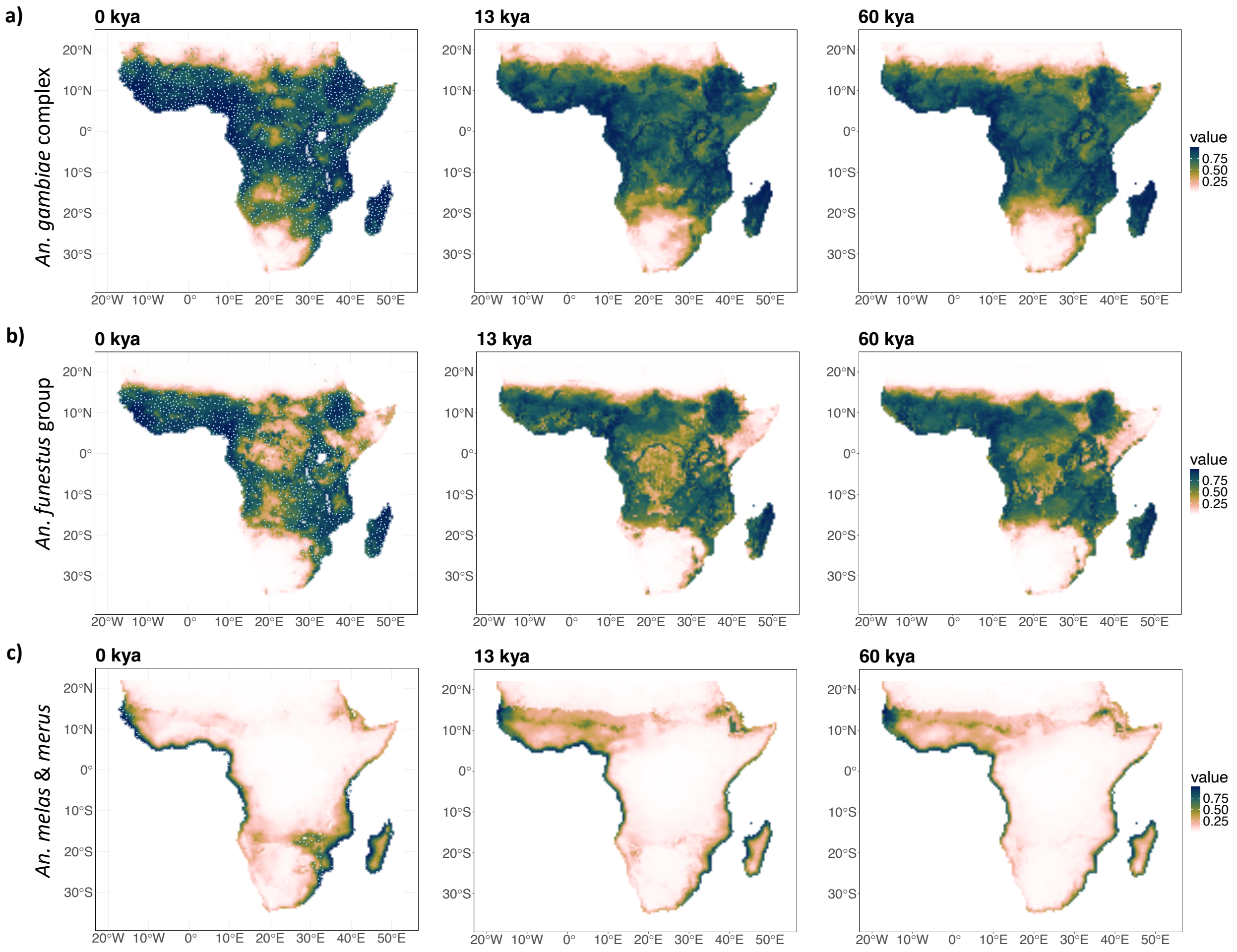


**Figure S3. Anopheles vectors SDMs for the present and the past.** These plots show the reconstructed niche of the studied vectors: a) *An. gambiae* complex at the present time (with observations included), at 13 and at 60 kya; b) *An. funestus* group at the present time (with observations included), at 13 and at 60 kya; and c) *An. melas* and An. merus at the present time (with observations included), at 13 and at 60 kya.

### Epidemiological information: the malaria stability index

The interplay between various climatic and epidemiological factors determines the relationship between vector and disease and the consequent variations in malaria transmission intensity. Therefore, we can consider the effects of multiple vectors on the potential risk of malaria (i.e., "stability" of malaria) by calculating a malaria stability index for each species. This can tell us what are the ecological conditions linked to high risk of malaria and, consequently, potential malaria's ranges.

Kiszewski and coworkers [54] suggest the contribution of mosquitoes as an objective measure of malaria transmission stability, which can be estimated through a spatial index. This index takes into account the characteristics of the vectors that are influenced by climate and that have an impact on vectorial capacity, such as mosquito survival rate, which affects the stability of transmission throughout the year, seasonal temperature and precipitation, which affects both mosquitoes and parasite life-cycle as well as extrinsic *P. falciparum* incubation duration.

Therefore, once the niche of dominant species of mosquitoes was identified through time, we calculated an index quantifying the potential risk of malaria at each time step, based on Kiszweski et al. (2004) [54]. The index includes across the months (m) the *P. falciparum* incubation period (E), proportion of biting people of the chosen dominant vectors (a), and daily survival rate (p) of the vectors:

$$\sum_{m=1}^{12} \boldsymbol{a_{i,m}^{2} p_{i,m}^{E} / -ln(p_{i,m})}$$

where $E = 111/Temperature - 16°C$ for *P. falciparum.*

Kiszewski et al. [54] report the human biting index (hbi) for various species that inform the proportion of biting people used in the equation.

To obtain the index, we firstly iterated through each time step (from 74 kya to present) obtaining the monthly temperature and precipitation variables. Each species SDM is considered separately, and we started by masking areas on the climate map where the species is not present to reduce computation over the areas that are excluded from analysis. Then the SDMs were filtered to exclude areas (or cells on the map) where the monthly temperature is under 15°C, and the precipitation of the previous month was lower than 10 mm as it is not viable for larvae. Then we can compute *E* (length of extrinsic incubation period in days) for the remaining cells. The maps for each month need to be multiplied for the SDM probabilities and then, to get a yearly measure, we sum the values across months. Finally, we take the maximum index value per cell across all species to calculate an overall stability index per year considering all vectors and, therefore, to find the areas with the highest risk rate.

In this way, the malaria stability index provides an indirect measure of the combined effects of the presence of multiple mosquito species at a given location on the potential risk of malaria as an infectious disease. Based on suitable climatic and environmental conditions and focusing on stability and transmissibility, this index expresses the potential stability of transmission of malaria [54], therefore, quantifying an overall potential risk of transmission. We note that a high stability index does not imply the presence of malaria, but rather defines its potential impact if it was present.

### Correlating the human niche to the malaria stability index

#### Summary of species distribution map generation for *H. sapiens*

Building on [74], we aimed to reconstruct the distribution of hunter-gatherer populations in Africa from 300 kya until the present. By modelling humans' evolutionary dynamics through a time-dependent SDM based on archaeological sites across the continent, we generated reconstructions of the human niche. Further details on methods can be found in [75].

**African hunter-gatherer archaeological sites**. The dataset included a total of 1242 sites from 120,000 cal.BP to 495 cal.BP [74,76] to form a curated dataset that would reflect the presence of hunter-gatherers in Africa. Therefore, only sites that were older than 8 kya and had no evidence of cultivation, pottery, or metal use [60,76,77] were included and filtered to obtain data points with published coordinates, radiometric dates, an age error range less than or equal to 20 ky (see [74] for more information on the quality control steps).

If 14C uncalibrated data were present, calibration was applied using the *rcarbon* R package [78] considering northern hemisphere archaeological localities (basic IntCal20 [79]) and southern hemisphere archaeological localities (basic SHCal20 calibration). Instead, the calibrated 14C ages were used to calculate the mean age (1σ error range). When an archaeological layer had both 14C and other dating methods, the 14C mean age and error were combined with those from the other methods to determine the final mean age for the models.

**Environmental and climatic data**. An SDM based on the curated dataset of archaeological sites was performed. The paleoclimatic reconstructions of interest [80] were accessed through the R package *pastclim* [81], including 17 BIOCLIM variables (excluding BIO2 and BIO3, as based only on daily summaries), net primary productivity (NPP) and leaf area index (LAI). The final model included five variables: leaf area index (LAI), temperature annual range (BIO7), mean temperature of the wettest quarter (BIO8), mean temperature of warmest quarter (BIO10), and precipitation of wettest quarter (BIO16). These variables were selected for their informativeness when comparing the habitat suitability for humans distribution in the *H. sapiens* presences against the distribution of 10000 randomly sampled points in the whole area [74].

**Model fitting**. Chronological uncertainty was respected by resampling 100 times each date from a truncated normal distribution defined by the mean and ±2σ [82].

The thinned dataset was used as presences (one presence for every 200 km radius was kept to avoid spatial autocorrelation, i.e., spatial thinning), and 200 pseudo-absence points were drawn for every presence. To ensure the climatic background was captured in our SDMs, each dataset was paired with 200 randomly sampled locations matched by time for each observation. This produced 100 datasets ("repeats") with varied sampled presences and background points, allowing repeated analyses to account for this stochastic sampling.

Then, a model ("time-varying-niche" model) was specified and its fit was evaluated using GAMs algorithm, using the R package *mgcv* [83]. In the model, the interactions between environmental variable and time (fitted as tensor products) were considered, and the GAM formula was included as:

```
gam(obs~ti(bio07, k=4) + ti(bio08, k=4) + ti(bio10, k=4)
        + ti(bio16, k=4) + ti(lai, k=4) + ti(time_bp, k=4)
        + ti(time_bp,bio07) + ti(time_bp,bio08) + ti(time_bp,bio10)
        + ti(time_bp,bio16) + ti(time_bp,lai),
        data=PA, family='binomial')
```

The following checks and limits were applied to all GAMs:

- a maximum threshold set to 4 for the degrees of freedom of the splines to avoid overfitting;
- checks on the residuals of the models such as Kolmogorov–Smirnov (KS) tests for correct distribution, dispersion and outliers (as in [74]), using the R package *DHARMa* [84];
- Akaike Information Criterion (AIC) was used for each repeat to verify that the time-varying niche models were best supported;
- Boyce Continuous Index (BCI) [85] was applied to assess the fit of the changing niche models, with an acceptance threshold of Pearson's correlation coefficient higher than 0.7 [86].

As the SDMs covered a period between 120 kya to 1 kya, projections were created to cover the time frame from 1 kya to the present. This was based on the projections for 105 kya, after testing the relationship between environment and time at 105, 110 and 120 kya.

The predictions of all repeats were averaged (mean and median) to create two ensembles, considering both AIC (for the time-varying niche) and BCI values above a threshold of 0.7.

For each repeat, the BCI was calculated by comparing ensemble predictions of the associated resampled presences against the predictions of the full set of resampled presences and background points. Then, the BCI for the ensemble is calculated as the mean/median over all repeats, with the mean retained for further analyses due to its better performance. The obtained probabilities of occurrences of the mean ensemble were converted into binary, considering only presence/absence. This was obtained by applying three thresholds and a modified version of the function *ecospat.mpa()* from the *ecospat* R package [87]: we identified the minimum predicted area encompassing 90% of our presences as "core area", a "peripheral area" (95% of presences) and a "total area" (99% of presences).

**Including the human niche**

The niche area was divided into core areas (the smallest area including 90% of archaeological sites/presences) and extended areas (covering 95% and 99% of sites respectively). We chose an intermediate level of 0.95 to include the extension of humans across the landscape.

Firstly, we checked the spatial overlay of the human core niche against the maps of malaria stability index extent across time (see Figure S4). Then, we compared the median of the index in the areas identified as core areas for humans against the median of the index in the areas outside the human range (i.e., areas identified as not suitable for human groups), considering upper and lower quantiles of 0.25 and 0.75. Desert areas (unsuitable for both humans and malaria vectors) were still included in this analysis.

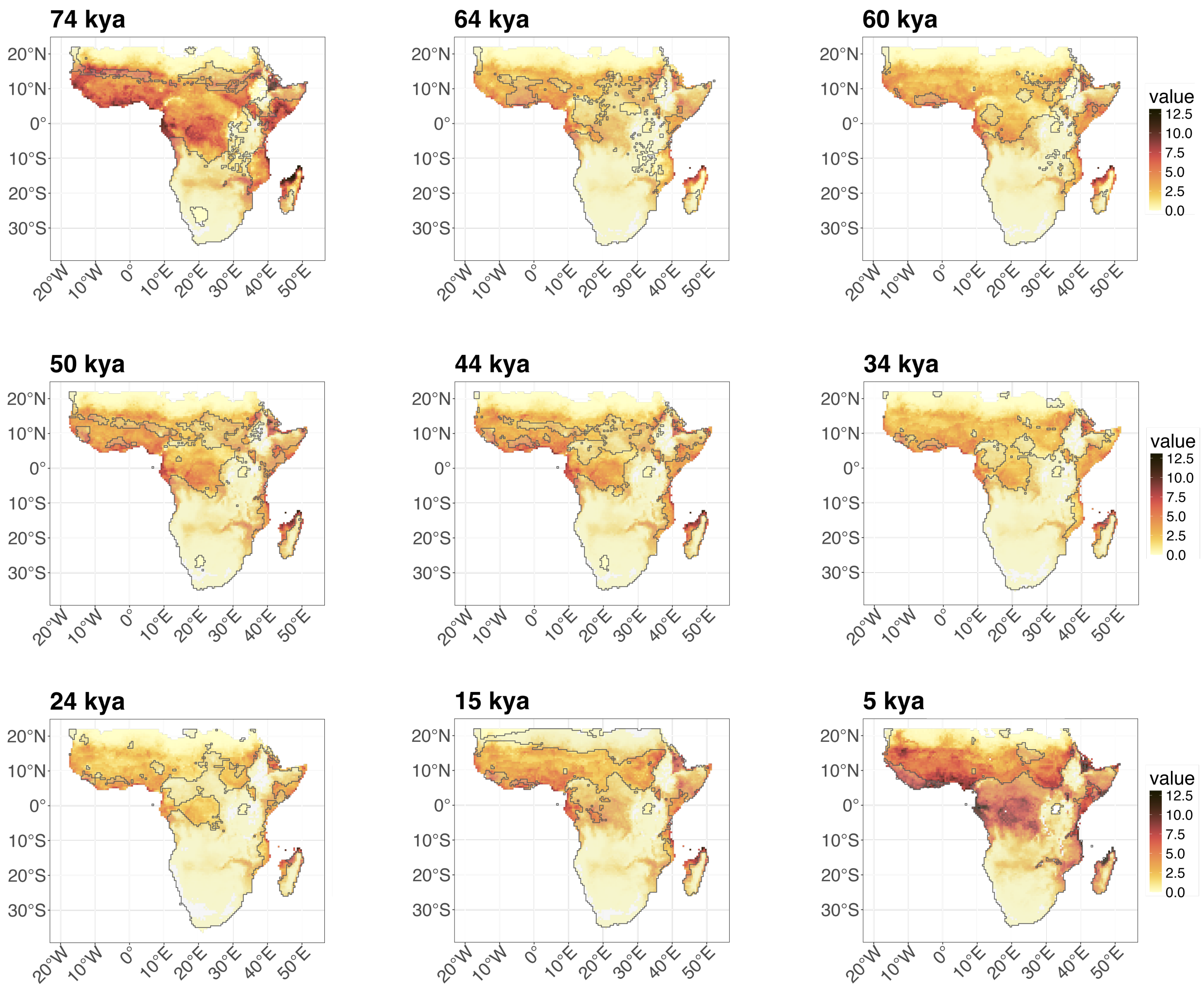


**Figure S4. Comparing the extent of human niche and malaria stability index through time.** These maps show the extent of the human niche (outlined in black) against the map of malaria stability index at nine time steps as an example: at 74 kya, 64 kya, 60 kya, 50 kya, 44 kya, 34 kya, 24 kya, 15 kya and 5 kya.

60 Yaw Asare, A., Mariangela, B. & Guiyun, Y. in *Current Topics in Malaria* (ed J. Rodriguez-Morales Alfonso) Ch. 20 (IntechOpen, 2016).

61 Snow, R. W., Amratia, P., Kabaria, C. W., Noor, A. M. & Marsh, K. The changing limits and incidence of malaria in Africa: 1939-2009. *Adv Parasitol* **78**, 169-262 (2012). https://doi.org/10.1016/b978-0-12-394303-3.00010-4
62 Beyer, R. M., Krapp, M. & Manica, A. High-resolution terrestrial climate, bioclimate and vegetation for the last 120,000 years. *Scientific Data* **7**, 236 (2020). https://doi.org/10.1038/s41597-020-0552-1
63 terra: Spatial Data Analysis v. R package version 1.7-8 (2024).
64 Goldewijk, K. K., Beusen, A., Doelman, J. & Stehfest, E. Anthropogenic land use estimates for the Holocene – HYDE 3.2. *Earth System Science Data* **9**, 927-953 (2017). https://doi.org/10.5194/essd-9-927-2017
65 Asner, G., Scurlock, J. & Hicke, J. Global synthesis of leaf area index observations: implications for ecological and remote sensing studies: Global leaf area index. *Global Ecology and Biogeography* **12**, 191-205 (2003). https://doi.org/10.1046/j.1466-822X.2003.00026.x
66 Ramankutty, N. & Foley, J. A. Characterizing Patterns of Global Land Use: An Analysis of Global Croplands Data. *Global Biogeochemical Cycles* **12**, 667-685 (1998). https://doi.org/http://dx.doi.org/10.1029/98GB02512
67 Elith, J. & Leathwick, J. R. Species distribution models: ecological explanation and prediction across space and time. *Annu. Rev. Ecol. Evol. Syst.* **40**, 677–697 (2009). https://doi.org/https://doi.org/10.1146/annurev.ecolsys.110308.120159
68 Leonardi, M., Colucci, M., Pozzi, A. V., Scerri, E. M. L. & Manica, A. tidysdm: Leveraging the flexibility of tidymodels for species distribution modelling in R. *Methods in Ecology and Evolution* (2024). https://doi.org/https://doi.org/10.1111/2041-210X.14406
69 Miller, E. F. *et al.* Post-glacial expansion dynamics, not refugial isolation, shaped the genetic structure of a migratory bird, the Yellow Warbler (Setophaga petechia). *bioRxiv* (2021). https://doi.org/2021.05.10.443405
70 Maxent software for modeling species niches and distributions (Version 3.4.1) v. 3.4.1 (2024).
71 Kuhn, M. & Wickham, H. *Tidymodels: a collection of packages for modeling and machine learning using tidyverse principles.*, <https://www.tidymodels.org/> (2023).
72 Roberts, D. R. *et al.* Cross-validation strategies for data with temporal, spatial, hierarchical, or phylogenetic structure. *Ecography* **40**, 913-929 (2017). https://doi.org/https://doi.org/10.1111/ecog.02881
73 Valavi, R., Elith, J., Lahoz-Monfort, J. J. & Guillera-Arroita, G. blockCV: An r package for generating spatially or environmentally separated folds for k-fold cross-validation of species distribution models. *Methods in Ecology and Evolution* **10**, 225–232 (2019). https://doi.org/https://doi.org/10.1111/2041-210X.13107
74 Hallett, E. Y. *et al.* Major expansion in the human niche preceded out of Africa dispersal. *Nature* (in press).
75 Padilla-Iglesias, C. *et al.* Pan-African metapopulation model explains Homo sapiens genetic and morphological evolution. *bioRxiv*, 2025.2005.2022.655514 (2025). https://doi.org/10.1101/2025.05.22.655514
76 Padilla-Iglesias, C., Grove, M. & Blinkhorn, J. Ecological drivers of hunter-gatherer lithic technology from the Middle and Later Stone Age in Central Africa. *Quaternary Science Reviews* **322**, 108390 (2023). https://doi.org/https://doi.org/10.1016/j.quascirev.2023.108390
77 Padilla-Iglesias, C. *et al.* Population interconnectivity over the past 120,000 years explains distribution and diversity of Central African hunter-gatherers. *Proc Natl Acad Sci U S A* **119**, e2113936119 (2022). https://doi.org/10.1073/pnas.2113936119

78 Crema, E. R. & Bevan, A. INFERENCE FROM LARGE SETS OF RADIOCARBON DATES: SOFTWARE AND METHODS. *Radiocarbon* **63**, 23-39 (2021). https://doi.org/10.1017/RDC.2020.95

79 Reimer, P. J. *et al.* The IntCal20 Northern Hemisphere Radiocarbon Age Calibration Curve (0–55 cal kBP). *Radiocarbon* **62**, 725-757 (2020). https://doi.org/10.1017/RDC.2020.41

80 Krapp, M., Beyer, R., Edmundsson, S. L., Valdes, P. J. & Manica, A. A comprehensive climate history of the last 800 thousand years. *EarthArXiv* (2019). https://doi.org/https://doi.org/10.31223/osf.io/d5hfx

81 Leonardi, M., Hallett, E. Y., Beyer, R. B., Krapp, M. & Manica, A. pastclim 1.2: an R package to easily access and use paleoclimatic reconstructions. *Ecography* **2023**, e06481 (2023). https://doi.org/https://doi.org/10.1111/ecog.06481

82 Will, M., Krapp, M., Stock, J. T. & Manica, A. Different environmental variables predict body and brain size evolution in Homo. *Nature Communications* **12**, 4116 (2021). https://doi.org/10.1038/s41467-021-24290-7

83 Wood, S. N. Fast stable restricted maximum likelihood and marginal likelihood estimation of semiparametric generalized linear models. *Journal of the Royal Statistical Society* **73**, 3-36 (2011).

84 DHARMa: Residual Diagnostics for Hierarchical (Multi-Level / Mixed) Regression Models. (2022).

85 Boyce, M. S., Vernier, P. R., Nielsen, S. E. & Schmiegelow, F. K. A. Evaluating resource selection functions. *Ecological Modelling* **157**, 281-300 (2002). https://doi.org/https://doi.org/10.1016/S0304-3800(02)00200-4

86 Hirzel, A. H., Le Lay, G., Helfer, V., Randin, C. & Guisan, A. Evaluating the ability of habitat suitability models to predict species presences. *Ecological Modelling* **199**, 142-152 (2006). https://doi.org/https://doi.org/10.1016/j.ecolmodel.2006.05.017

87 Di Cola, V. *et al.* ecospat: an R package to support spatial analyses and modeling of species niches and distributions. *Ecography* **40**, 774-787 (2017). https://doi.org/https://doi.org/10.1111/ecog.02671
